# Reconstitution of Helical Soluble α-Synuclein through Transient Interaction with Lipid Interfaces

**DOI:** 10.1101/202994

**Authors:** Matteo Rovere, John B. Sanderson, Luis Fonseca-Ornelas, Tim Bartels

## Abstract

α-synuclein (αSyn) is one of the key players in the pathogenesis of Parkinson’s disease (PD) and other synucleinopathies. Its misfolding and subsequent aggregation into intracellular inclusions are the pathological hallmark of these diseases and may also play a central role in the molecular cascade leading to neurodegeneration. In this work, we report the existence of a novel soluble α-helical conformer of αSyn, an archetypal “intrinsically disordered protein” (IDP), obtained through transient interaction with lipid interfaces. We describe how the stability of this conformer is highly dependent on the continuous, dynamic oligomerization of the folded species. The conformational space of αSyn appears to be highly context-dependent, and lipid bilayers might play crucial roles as molecular chaperones for cytosolic species in a cellular environment, as they do in the case of this previously unreported structure.

**Significance Statement:** Both genetic and histopathologic evidence tie α-synuclein (αSyn) to the pathogenesis of Parkinson’s disease (PD), a widespread neurodegenerative disorder. Lipids play a central role in the dynamics of αSyn in physiology and disease. αSyn undergoes a coil-to-helix transition when binding to lipid vesicles and it is involved in the regulation of synaptic vesicle trafficking. Furthermore, recently discovered α-helical, aggregation-resistant “multimers” of αSyn could constitute a protective conformational pathway. We report the existence of a folded, lipid-unbound αSyn conformer that forms upon transient interaction with lipids and is stabilized by dynamic homooligomerization, suggesting that synaptic activity could modulate resistance towards aggregation. Our results are therefore important both for the molecular pathology of PD and the structural biology of intrinsically disordered proteins.

## Introduction

Parkinson’s disease (PD) is the second most common neurodegenerative disorder after Alzheimer’s disease with a prevalence of approximately 1% in people over 60 years of age, increasing to 4% by age 85 (1). Among other risk and causative factors, both genetic and histopathologic evidence convincingly implicate α-synuclein (αSyn) in the pathogenesis of PD. SNCA (PARK1), which encodes αSyn, was the first gene to be linked to PD (2) and, since then, both point mutations and duplications/triplications of the locus have been shown to cause PD with a Mendelian pattern of inheritance (3, 4). In addition, the neuropathological hallmarks of PD and other so-called synucleinopathies (5) are intracellular protein aggregates known as Lewy bodies and Lewy neurites that present a fibrillar, β-sheet-rich core of αSyn (6).

αSyn is a small, 140 amino acid-long, protein of about 14 kDa that belongs to a family of proteins including β- and γ-synuclein. Its highly conserved N-terminus contains 7 repeats of an 11-residue imperfect XKTKEGVXXXX motif similar to those present in the lipid-binding amphipathic domains of class A2 apolipoproteins (7).

Lipids play a central role in αSyn’s dynamics in physiology and disease. Despite its classification as an archetypal “intrinsically disordered protein” (IDP) (8, 9), αSyn adopts a predominantly α-helical structure with a disordered C-terminus when bound to lipid interfaces (10, 11). Its affinity is highest for highly curved bilayers of negatively charged lipids, similar in both radius and lipid composition to synaptic vesicles (10). Its exact physiological function remains unknown, but it has been increasingly associated with the regulation of synaptic vesicle exocytosis and recycling, suggesting that its dysfunction may cause the defective dopaminergic neurotransmission found in PD (12–17). Several works on the pathobiology of synucleinopathies have also shown that αSyn overexpression and familial-PD-associated mutants cause trafficking defects in various model organisms by interfering with the docking and fusion processes of vesicles (18–20). Furthermore, mutations in the β-glucocerebrosidase gene (GBA) represent a significant risk factor for PD. GBA mutations are associated with Gaucher’s disease, an autosomal recessive lysosomal storage disorder caused by the dysfunctional metabolism of sphingolipids, with Parkinsonism among its neurological symptoms (21). Heterozygous carriers of GBA mutations have a five-fold increase in the risk of developing PD (22).

Recently, the purification and characterization of a soluble, α-helical, tetrameric αSyn assembly under non-denaturing conditions challenged the classic paradigm of its natively unfolded nature (23, 24). Though controversial at first (25, 26), our group and others have since reproduced and extended these findings (27–29). The discrepancy between the classic “natively unfolded” characterization of αSyn, obtained from denaturing purifications and *in vitro* studies of recombinant protein (8), and the new evidence of mammalian-cell-isolated, α-helical αSyn, is believed to be linked to the absence of specific eukaryotic co-factors and chaperones (28, 30).

These results underscore the importance of characterizing the conformational space of disordered proteins, particularly αSyn, in the presence of cytosolic folding determinants such as molecular crowding, interacting proteins, and lipid bilayers. Given the well-characterized propensity of αSyn to form α-helical-rich structure upon interaction with vesicles and the evidence supporting the existence of cytosolic α-helical “multimers” (31) of αSyn, it is reasonable to imagine a role for lipids in this multimerization process. In this work, we investigate how lipid interfaces can assist the refolding of a α-helical, soluble conformer of αSyn through transient interaction. In order to do so, we use the phase transition of phosphatidylcholine (PC) small unilamellar vesicles (SUVs) to modulate αSyn’s lipid binding affinity and characterize the properties and the stability of the conformer thus obtained.

## Results

### Temperature-dependent αSyn-lipid interaction shows hysteresis in α-helical fold

αSyn has been extensively shown to have a central role in defining the organization and the ultrastructure of the synaptic terminus (17, 32). The membrane-remodeling effect of its bound α-helical conformation (27, 32, 33) is also a highly dynamic process, depending on a variety of complex factors as membrane curvature, electrical activity and membrane microheterogeneity, especially in the forms of lipid rafts (34, 35). To mimic the finely tunable lipid affinity and conformational flexibility of αSyn in a cellular setting, we used the binding of αSyn to PC SUVs undergoing a phase transition (36), which has been shown to be the mechanism underlying the localization of αSyn to lipid rafts (34, 37). The affinity of αSyn for zwitterionic PC SUVs is driven by the insertion of αSyn in the lipid packing defects, rather than by the electrostatic interactions involved in its well-characterized affinity for anionic vesicles (10) and thus, is temperature-dependent (36, 38). Below their melting temperature (T_m_), SUVs are highly strained gel-like (minimally fluid) systems with low curvature radii that cause an imperfect distribution of PC molecules, which allows αSyn to insert and improve the organization of the double layer. The protein folds in a predominantly α-helical structure that enters the hydrophobic core of the liposome and remodels the lipid environment. Since the formation of packing defects induced by the liquid-disordered to gel-state phase transition triggers this event, αSyn binding and folding do not take place above the T_m_ of the respective lipids (36). This unique behavior, coupled with the choice of an adequate acyl chain length, can be exploited to control the binding and folding of the protein by variation of the sample temperature within the range of stability of secondary structures. In our protocol, we first mixed SUVs and αSyn well above the T_m_ and then lowered the temperature to allow membrane binding and formation of the protein’s α-helical secondary structure. Heating the sample above the T_m_ should then release αSyn from the lipid vesicle, potentially preserving its secondary structure (Fig. 1A). These events can be easily monitored using the circular dichroism (CD) spectrum of the protein to quantify the helix-coil conformational change accompanying membrane interaction (10). As shown in Fig. 1B, controlled melting of the lipid double layer preserved helical fold in the protein and allowed further characterization of the conformers of αSyn reconstituted in the process. αSyn can retain the α-helical fold, measured using the signal of the minimum at 222 nm, at much higher temperatures in the upscan branch of the curves than during the downscan, which shows the binding taking place roughly at the T_m_ of the SUVs (~14°C) (Fig. 1C). The degree of hysteresis presents a strong negative correlation with the temperature gradient speed used in the melting curve. While moving from 0.2 to 0.5°C/min. doesn’t change significantly the outcome, increasing the ramp to 1.0 or 2.0°C/min. leads to the loss of most of the fold at RT (Fig. 1C), though some α-helical content remains when compared to the downscan signal. With the increase in the temperature gradient steepness, the oscillations in the signal around the T_m_ increase in amplitude as well, possibly due to critical fluctuations originating from αSyn-induced membrane microdomains (39, 40).

**Fig. 1.**
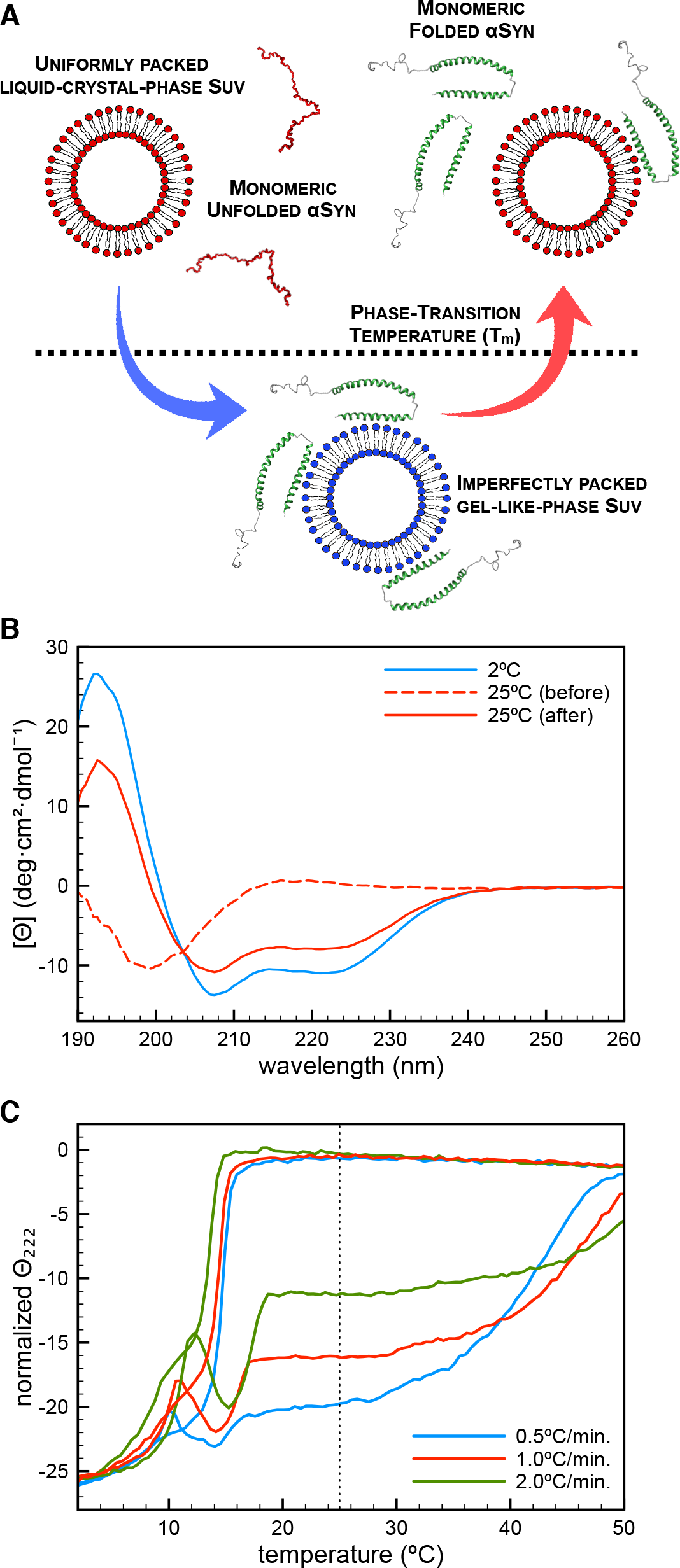
The temperature-dependent binding behavior of αSyn leads to hysteresis in α-helical content. (A) Diagram of the protocol developed for the preparation and isolation of folded αSyn intermediates. PC SUVs, freshly prepared, were mixed with recombinant αSyn (1:1000-1200 protein:lipid molar ratio) and cooled below their phase transition temperature (T_m_), then heated at an equal rate to above their T_m_, ultracentrifuged at 170,000xg for 1 hr., and analyzed. (B) Circular dichroism (CD) spectra of a sample of αSyn refolded via temperature cycling in the presence of 13:0 PC SUVs, measured at 25°C *(after)*. The CD spectra of the sample at the lower end of the temperature scan *(2°C)* and before the temperature cycling protocol *(before)* are shown for comparison. (C) Ellipticity signal at 222 nm of 5 μM αSyn refolded via temperature cycling in the presence of 13:0 PC SUVs, followed via circular dichroism (CD) readings recorded every 0.5°C between 2°C and 50°C, varying the temperature gradient steepness. The dotted line indicates RT (25°C).

In order to test whether the results originated from the residual affinity of αSyn for lipid membranes at temperatures higher than their T_m_, we studied the binding of αSyn to 13:0 PC SUVs using isothermal titration calorimetry (ITC). Binding sigmoids recorded at different temperatures highlight the sharp decrease in the protein-lipid affinity following the increase in temperature (Fig. 2), showing that no binding to the SUVs can be detected not only at T>T_m_, but also at temperatures close to the phase transition. Since the 222-nm CD signal for αSyn titrated with 13:0 PC SUVs plateaus at a protein:lipid ratio of about 1:1000 (Fig. S1), we used protein:lipid ratios ranging from 1:1000 to 1:1200 for all our experiments. Temperature-cycling experiments were also performed using three other PC lipids with saturated acyl side chains (12:0, 14:0 and 15:0 PC), in order to determine whether the hysteresis observed when recording melting curves in the presence of 13:0 PC SUVs was specific for this acyl chain length or common to related phosphatidylcholines. Melting curves showed that, although the phenomenon of hysteresis in the α-helical CD signal seems to be common to all of the saturated phospholipids screened, the shape of the melting curve depends on the acyl chain length (and thus on the T_m_ of the lipids). The highest fold retention can be achieved with 13:0 PC SUVs (Fig. S2), which have a T_m_ of about 14°C (Fig. 3A). This specificity does not seem to depend on the varying affinity (measured using isothermal titration calorimetry, ITC) of αSyn for the different lipid assemblies (Fig. S3) or the size of the SUVs (Supporting Table 1), which show no clear correlation with the αSyn-lipid behavior. Finally, it must be noted that N-alpha acetylation, a post-translational modification shown to be characteristic of the near totality of physiological αSyn (23, 41) appears to affect the hysteresis process favoring the helical form of αSyn (Fig. S4), thus providing additional proof of its importance in lipid binding and helical multimer formation (42, 43).

**Fig. 2.**
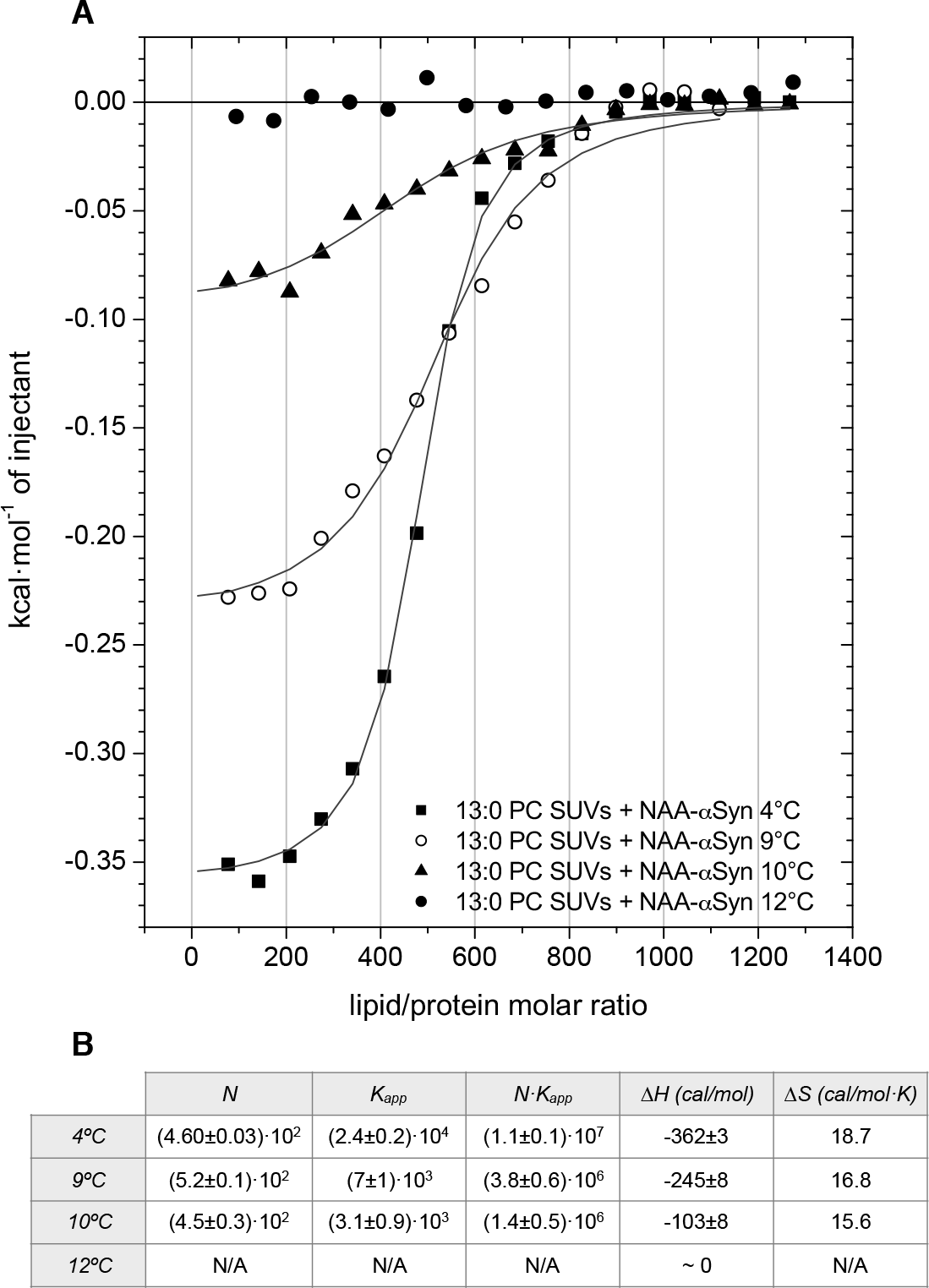
αSyn’s membrane-binding is abolished as the temperature approaches T_m_. (A) Isothermal titration calorimetry (ITC) binding curves of 5μM αSyn titrated with freshly prepared 13:0 PC SUVs, plotted with their fitting function curves (solid lines). After integration of the differential heat signal, a N independent binding sites model was used to fit the data. Experiments were performed at varying temperatures from 4°C to 12°C. (B) Thermodynamic and stoichiometric parameters obtained from the fitting of the binding curves, along with standard errors.

**Fig. 3.**
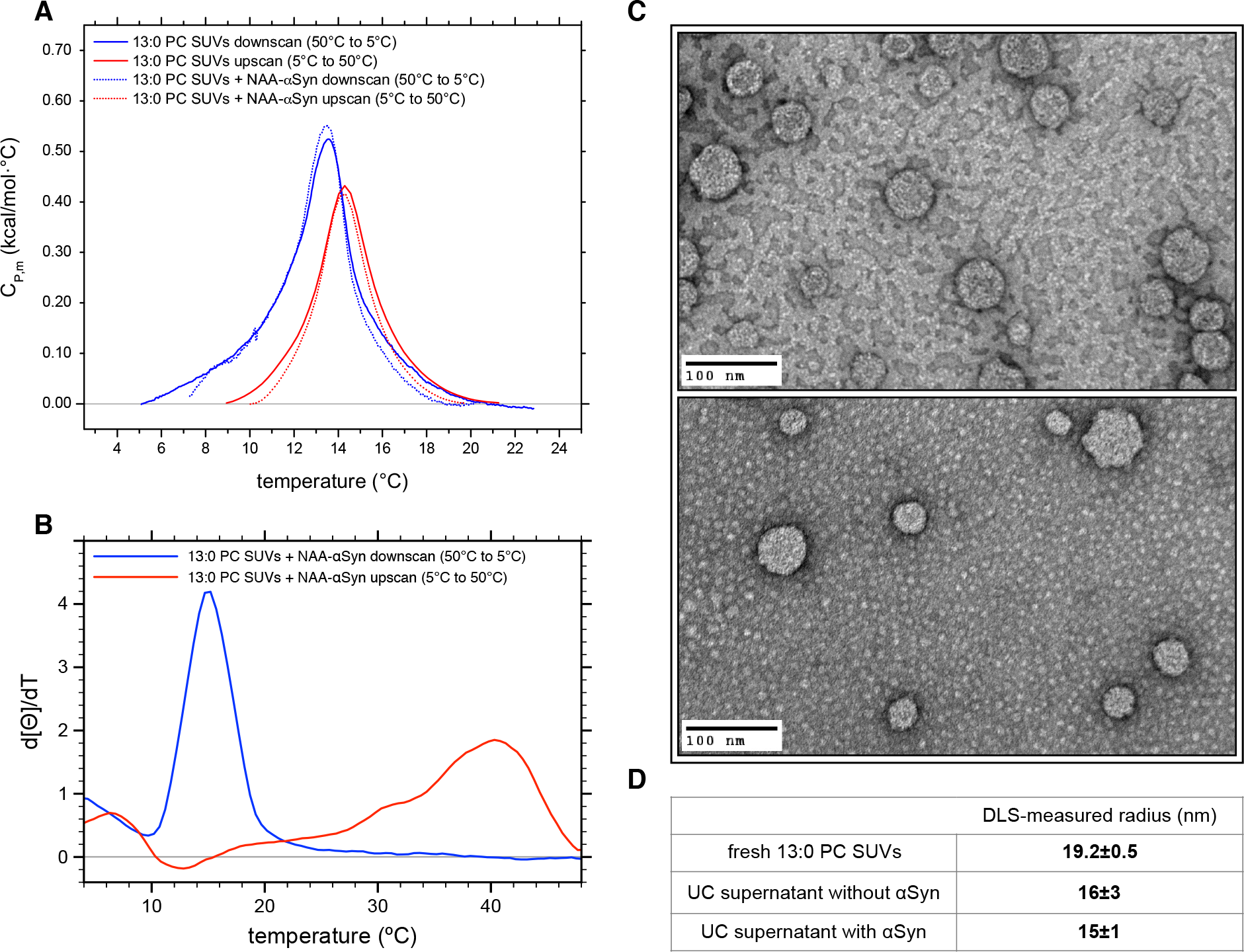
The fold hysteresis of αSyn is uncoupled from the SUVs’ phase transition. (A) Normalized differential scanning calorimetry (DSC) thermograms of 2.75 mM 13:0 PC SUVs either alone (solid curves) or in the presence of 2.5 μM αSyn (dashed curves). Temperature scans were performed using a 0.5°C/min. (30°C/hr.) ramp between 50°C and 5°C (50°C-5°C-50°C). The blue curves indicate downscans (50°C-5°C), while the red ones indicate upscans (5°C-50°C). Linear baselines were subtracted from the peaks and no other transitions were recorded in the 5-50°C temperature range. (B) Numeric first derivative of a smoothed 222-nm ellipticity signal, followed via circular dichroism (CD), of 5 μM αSyn refolded in the presence of 13:0 PC SUVs through a temperature scan between 2°C and 50°C (50°C-2°C-50°C) at 0.5°C/min., recording the signal every 0.5°C. The blue curve shows the 50°C-2°C downscan, while the red one shows the upscan (2°C-50°C). (C) Transmission electron microscopy (TEM) micrographs of the supernatant fraction of 12 mM 13:0 PC SUVs subjected to temperature cycling (50°C-2°C-25°C) alone *(top)* or in the presence of 10 μM αSyn *(bottom)* and collected after a 170,000xg ultracentrifuge spin. Samples were negatively stained with 1% uranyl formate (UF). Micrographs shown are taken at 65,000x magnification. (D) Mean radii of liposomes, measured by dynamic light scattering (DLS) at 25°C, found in the supernatant fraction of 12 mM 13:0 PC SUVs subjected to temperature cycling (50°C-2°C-25°C) alone *(without αSyn)* or in the presence of 10 μM αSyn *(with αSyn)* and collected after a 170,000xg ultracentrifuge spin. Error bars indicate SDs obtained from 3 independent experiments. The mean radius and SD (N=3) of a dispersion of fresh 13:0 PC SUVs is shown for comparison.

### αSyn’s conformational hysteresis is uncoupled from the phase transition of the SUVs

ITC can only measure lipid association under isothermal conditions. However, the hysteresis in αSyn’s helical content could be just paralleling a structural hysteresis in the lipid bilayers, determining the release of the protein at a much higher temperature than the T_m_ during the upscan to 50°C. In order to examine the extent of the dynamic interplay between αSyn and the SUVs during (and after) the phase transition, we used differential scanning calorimetry (DSC) to look for a link between protein fold hysteresis and the transition temperature of the liposomes. It has been reported that the presence of αSyn can markedly reshape the phase transition, not only by shifting the T_m_ (36, 44), but also by changing the heat profile of the DSC curve, signalling an increase in the cooperativity of the process (36). We thus recorded both downscan and upscan heat capacity curves of 13:0 PC SUVs both with and without αSyn, using the same parameters as for the CD temperature scans. In the lipid-only scan, just one sharp peak is detected (both in the downscan and the upscan branch of the curve), corresponding to the phase transitions of the vesicles (Fig. 3A). The T_m_ shows a minor hysteresis (13.5°C in the downscan, 14.3°C in the upscan), a commonly observed phenomenon in lipid transitions, even when using slow ramps such as the one employed in these scans (30°C/hr.) (45). The DSC curves recorded in the presence of the protein are virtually superimposable to the ones recorded in its absence (Fig. 3A). This implies that any effect on the lipid behavior due to bound protein must be extremely small and undetectable as an enthalpy change, as the areas of the peaks recorded in the experiments are the same and αSyn’s concentration was too low to provide a detectable heat signal. This strongly argues against the possibility that αSyn’s conformational hysteresis is due to a “belated” phase transition of the SUVs.

To further compare the lipid behavior during temperature cycling to the conformational transitions of the protein, we took the first derivative of the CD signal of a representative 13:0 PC SUVs-αSyn scan. The differentiated temperature scan curve shows an excellent accord with the DSC in the downscan branch, showing just one sharp peak at roughly the same temperature as the T_m_ of the lipids (14.9°C) (Fig. 3B). On the other hand, there is minimal change in ellipticity around the T_m_ in the upscan, with only one broad peak around 40°C (accompanied by a shoulder around 30°C), indicating the unfolding of the protein in the absence of structural changes in the lipid membrane.

Finally, we studied the morphology and size of the lipid assemblies after a temperature scan, performed both in the presence and absence of αSyn, to check for possible remodeling effects of αSyn on the SUVs (33, 46). Supernatants were freshly collected and analyzed after the ultracentrifuge spin performed at the end of the temperature cycling, done to remove larger lipid aggregates formed by the fusion of PC SUVs below their T_m_ (47). Transmission electron microscopy (TEM) micrographs show no difference in shape or size between the samples (Fig. 3C) and no difference in the hydrodynamic radii measured by dynamic light scattering (DLS) was detected (Fig. 3D), further indicating that the structural changes in the lipid bilayers are completely reversible and cannot explain the conformational hysteresis of αSyn.

### Lipid interfaces act as molecular chaperones for soluble αSyn

To confirm that the α-helical conformer of αSyn existed in a soluble, not lipid-bound, state and quantify the relative amounts of the two populations, we used both size-exclusion chromatography (SEC) (along with an ELISA-based quantification of αSyn and phosphate analysis in order to measure the 13:0 PC content in the SEC fractions) and TEM imaging coupled with immunogold labeling of αSyn (Fig. 4A,B,C). For SEC, the supernatant from an ultracentrifuge spin of a refolded αSyn-lipid mixture was immediately loaded on a Superdex 200 10/300 GL column and 1 mL fractions were collected and analyzed as described. The 280 nm absorbance profile in the chromatogram shows three peaks (Fig. S5): the third and last peak, undetectable both in the ELISA and in the phosphate analysis (Fig. 4A), likely indicates the elution of small lipid assemblies, running close to the ions peak seen in the conductivity profile. The first peak, running roughly at the column’s void volume (MW>600kDa), corresponds to the intact SUVs, as confirmed by the phosphate analysis indicating the presence of lipids only in the first peak fractions (Fig. 4A). The majority of αSyn (~80%) elutes, free from lipids, in the second peak, around 15 mL. This indicates a MW of about 60 kDa, in accordance with the extended conformation of αSyn (8). A fraction of the population of αSyn (~20%) co-elutes with the SUVs in the first peak, indicating the presence of a small amount of αSyn still bound to the membranes. The percentage of lipid-bound and lipid-unbound protein was obtained from the quantification by ELISA of the areas of the two chromatographic peaks from repeated experiments (phosphate analysis confirmed the absence of lipids in the second, membrane-unbound, protein peak) and compared to the percentage of folded protein after a temperature scan. If only the membrane-bound population of αSyn were α-helical, we would see an approximate drop of 80% in the helicity of our samples, while we only observe a decrease of 30-35% in the percentage of helical fold. Using another independent technique, TEM imaging coupled with immunogold labeling of αSyn, we repeated the quantification of the two protein populations (lipid-bound and unbound) and once again found that αSyn is largely unbound from the SUVs (~80%, in excellent accordance with the SEC/ELISA data) (Fig. 4B,C).

**Fig. 4.**
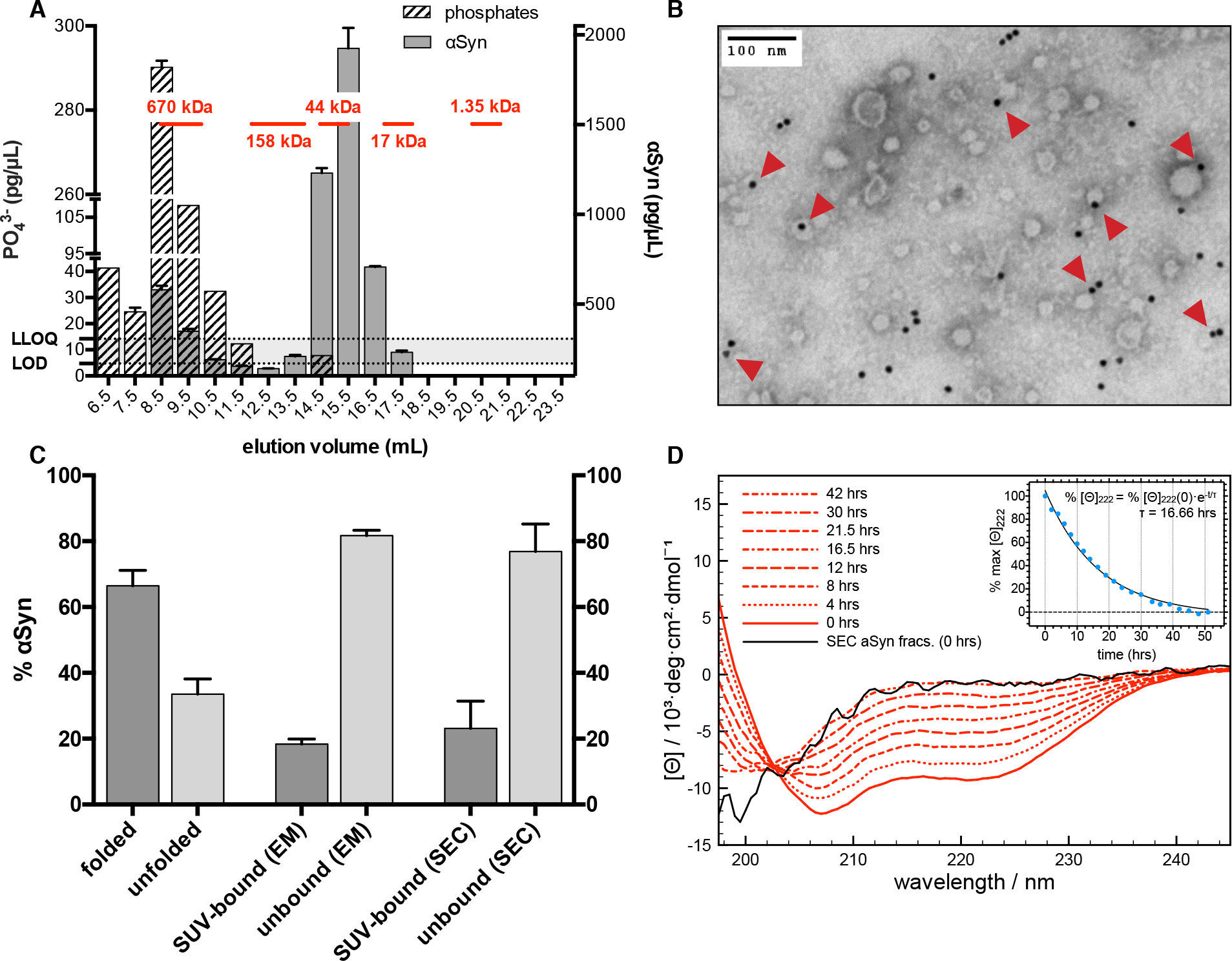
α-helical αSyn is largely unbound from the SUVs and exists as a metastable conformer. (A) ELISA-measured αSyn concentration and phosphate concentration, obtained from quantitative phosphate analysis, in the SEC fractions of a refolded αSyn sample run on a Superdex 200 column. Bars are labeled with their mid-fraction elution volume. Black lines indicate the approximate peak widths of the SEC standards run on the column, along with their molecular weights. 1-mL fractions were collected between 6 mL and 24 mL. SDs are plotted for each fraction, obtained from technical duplicates. The limit of detection (LOD) and the lower limit of quantification (LLOQ) for the phosphate analysis are indicated with dotted lines. (B) Representative transmission electron microscopy (TEM) micrograph of an αSyn sample refolded via temperature cycling in the presence of 13:0 PC SUVs (30,000x magnification). The sample was negatively stained with 1% phosphotungstic acid (PTA) and immunogold-labeled for αSyn using the 15G7 antibody. Red arrows indicate gold-nanoparticle-conjugated αSyn molecules counted as bound to SUVs. (C) Bar diagram comparing the percentage helical fold retention of αSyn (quantified via the 222nm signal at 25°C obtained from renormalized CD spectra measured after ultracentrifugation in 3 independent experiments) with the percentage of αSyn still bound to the SUVs quantified by two independent techniques, SEC (N=5) and EM (N=10 micrographs from 3 independent experiments, a total of 1292 immunogold particles and 2256 SUVs were counted). αSyn concentration in SEC fractions was quantified via ELISA and fractions were defined as lipid-bound or lipid-unbound using quantitative phosphate analysis. (D) Circular dichroism (CD) spectra of αSyn refolded via temperature cycling in the presence of 13:0 PC SUVs and measured at 25°C over 42 hrs. The CD spectrum of the lipid-unbound αSyn peak fractions from an SEC run of a sample of temperature-cycled αSyn is also shown (“SEC αSyn fracs”). The peak fractions were concentrated at RT with a spin filter and measured at 25°C immediately following elution from the SEC column. (inset) Percentage of the 222-nm molar ellipticity signal (100% at 0h). The data were fitted with an exponential curve, with % [Θ]_222_(0) and the decay time (t) as free parameters and with the constraint that the absolute value of the plateau should be smaller than % [Θ]_222_(42 hrs).

We then moved on to characterize the stability of the soluble helical αSyn conformer. The slow temperature ramp required to maintain most of the α-helicity already suggested the presence of an irreversible process and, by extension, the existence of a kinetically trapped conformer (Fig. 1C). A time course measurement, monitoring the helical content of αSyn, confirmed the metastable nature of the protein structure reconstituted using our protocol. While the CD spectrum initially shows prominent α-helicity, the folded content disappears in an exponential fashion over the course of 42 hrs (Fig. 4D).

### Stabilization of helical αSyn arises from dynamic multimerization

In view of the past reports on helical αSyn multimers, both *in vitro* and *in vivo* (23, 28, 48), we set to study the oligomerization state of the helical conformers of αSyn using disuccinimidyl glutarate (DSG) cross-linking (28). Samples from the supernatant, kept at RT, were collected immediately following the UC spin, and after 18 hrs or 36 hrs, and cross-linked with 500 μM DSG. Western blotting shows the cross-linked samples along with negative controls of recombinant αSyn cross-linked at the same concentration (of protein and DSG) and a positive control of intact M17D cells stably expressing wt αSyn to show the characteristic “physiologic” pattern of multimeric bands (28) (Fig. S6). The band pattern in each of the supernatant lanes resembles that of recombinant monomeric αSyn in the absence of lipids, indicating that the protein does not form stable, defined multimers as the ones previously characterized (23, 28), but could well be involved in a dynamic multimerization process that stabilizes its folded form (24). This possibility is consistent with the finding that a CD measurement of the unbound αSyn fractions, concentrated immediately after SEC fractionation, returns the spectrum of an unfolded protein (Fig. 4D). If a highly dynamic multimer formation was the cause of the long lifetime of folded monomeric αSyn, a drastic dilution as the one underwent on the SEC column (about 1:80) would markedly reduce the rate of these multimolecular processes leaving the irreversible unfolding as the dominant event, thus speeding up the decay in the CD signal and making it impossible to directly measure the unfolding kinetic of SEC-isolated αSyn.

Since the techniques employed up to this point were limited in their ability to detect highly dynamic populations of multimers, we used NMR spectroscopy in an attempt to answer this question (Fig. 5A). For this purpose we could only rely on the resonances in the aromatic/amidic region, since unfortunately the protein signal in the rest of the spectrum is masked by that of the free 13:0 PC. The severely broadened and barely detectable signal of lipid-bound αSyn, in the presence of 13:0 PC SUVs below their T_m_, is also shown for comparison. Inspection of the peak pattern and the linewidths of the refolded sample (compared with the spectrum of recombinant, unfolded protein at the same temperature) shows that the protein is unbound from lipid interfaces and closely replicates the peak profile of soluble αSyn. The slight peak broadening and lower intensities suggest a slower tumbling rate of the species observed, consistent with the multimerization of refolded αSyn but also, for example, with a binding/unbinding membrane exchange (49).

**Fig. 5.**
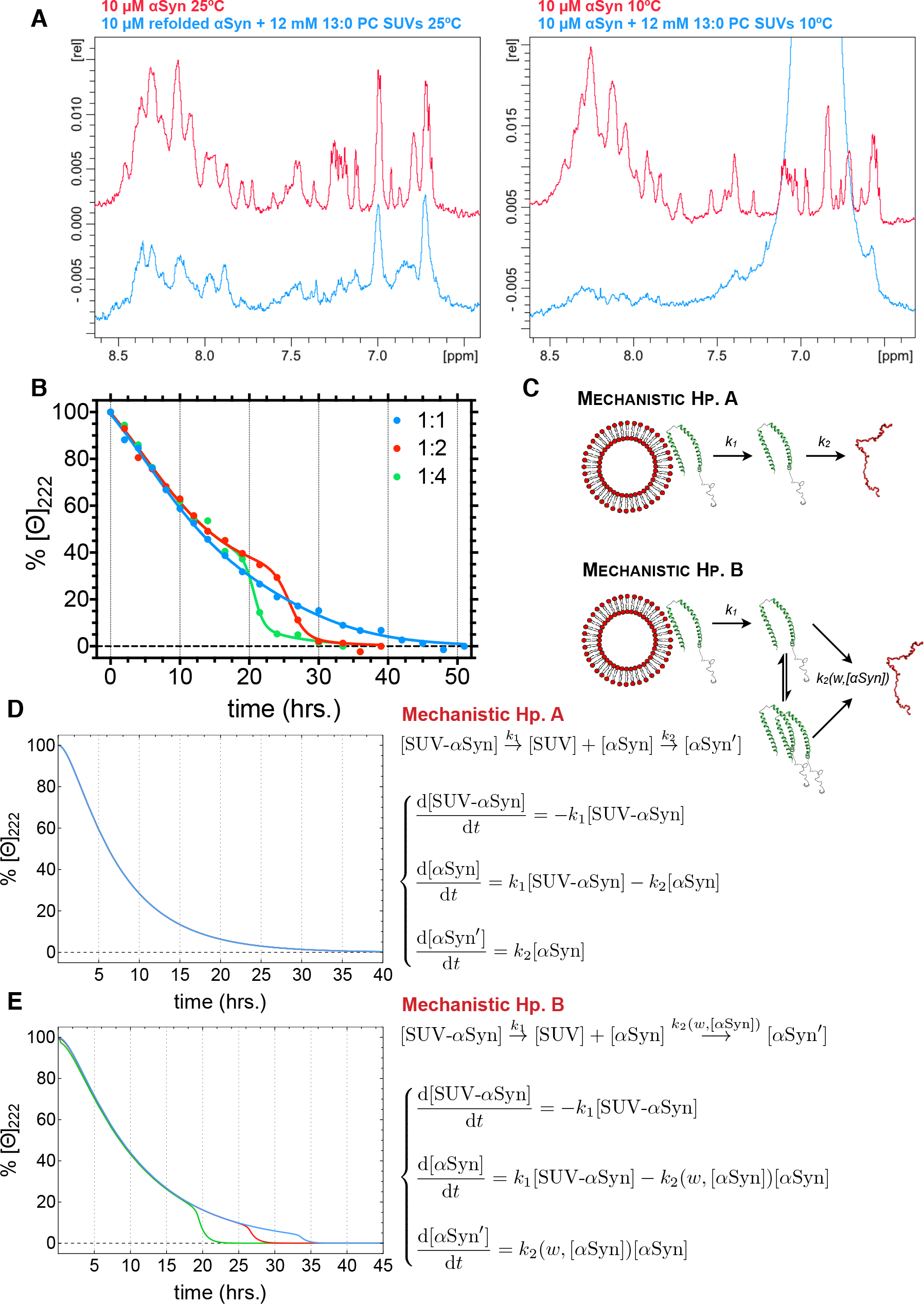
αSyn’s unfolding curves indicate a complex mechanism involving dynamic multimerization. (A) ^1^H-NMR spectrum of 10 μM αSyn in the presence *(blue)* or absence *(red)* of 12 mM 13:0 PC SUVs, measured at either 25°C *(left)* or 10°C *(right).* Only the aromatic/amidic region of the NMR spectrum of the protein is shown, as elsewhere the signal of free lipids masks that of the protein (one lipid peak also overshadows part of the protein signal at 10°C). The signal of the bound protein is severely broadened and barely detectable, whereas refolded αSyn closely reproduces the peak pattern of the unbound protein while showing a certain degree of broadening consistent with our mechanistic hypothesis. (B) Percentage of the 222-nm molar ellipticity signal (set at 100% at 0h), obtained from CD measurements of a sample of αSyn refolded via temperature cycling in the presence of 13:0 PC SUVs (1:1) and its dilutions in 10 mM (NH_4_)Ac, pH 7.40 (1:2 and 1:4). % [Θ]_222_ values are shown at time points following the ultracentrifuge spin. Global fitting of the data using a biphasic (double-sigmoid) model was performed sharing all but two parameters among the curves and setting the plateau at 0% (See Fig. S7 for details). (C) Diagram of the two mechanisms tested with the kinetic analysis of the CD-followed unfolding of αSyn. Hp. A is the classical view of αSyn’s interaction with membranes, seeing the binding to the interfaces as a critical and irreversible step in the coil-to-helix transition of the protein. Through the inclusion of a dynamic multimerization step, Hp. B accounts for the shape of the measured curves and the changes observed following serial dilutions of the sample, and is thus the proposed mechanism. (D) and (E) show the simulated curves (calculated for the three dilution factors measured) and the kinetic equations corresponding, respectively, to Hp. A and B.

To exclude this possibility and further validate our hypothesis that the metastability of the folded αSyn conformer depends on intermittent multimerization, we measured via CD the unfolding kinetics of an undiluted refolded αSyn sample and of 1:2 and 1:4 dilutions of the same sample in buffer (Fig. 5B). A CD spectrum of each of the three samples, incubated at RT, was obtained every 2-3 hrs until four superimposable spectra in a row were recorded, indicating the endpoint of the unfolding process. The percentage of the molar ellipticity at 222 nm was used to follow the loss in secondary structure and the transition to random coil. We see a marked decrease in the time at the endpoint moving from the 1:1 sample (51 hrs) to the ones diluted 1:2 (39 hrs) and 1:4 (33.5 hrs), which would not be the case if the process were one of concerted unbinding and unfolding from a vesicle-bound state (Fig. 5C). On the other hand, this observation is consistent with a multimolecular event as the transient homooligomerization process, which would be slowed down by dilution (Fig. 5C). Decreasing the concentration of soluble, folded αSyn would decrease the rate of the self-association as well, causing a faster loss of fold of the helical monomer. Given the complex, biphasic shape of the unfolding curves, an attempt to develop a mechanistic hypothesis for the observed trend was made. The classic hypothesis regarding αSyn’s folding upon contact with membranes was compared to our new paradigm postulating a wider range of stability for the folded protein, also existing as a long-lived conformer when unbound from lipid interfaces (Fig. 5C). Kinetic equations for the unbinding and unfolding from lipid bilayers were drafted, solved analytically and plotted (Fig. 5D), showing no dependence of the unfolding profile from the dilution factor (as intuitively expected, since such a mechanism would not involve any multimolecular steps, see SI Appendix for details). In addition to confirming that the unfolding of αSyn cannot be attributed to a slow unbinding of the protein from membranes, the failure of this model to reproduce experimental data also displays the necessity to introduce a new multimolecular event accounting for the sigmoidal unfolding profile clearly observed in the curves measured from samples diluted 1:2 and 1:4 (and most likely present in the 1:1 curve as well, but almost undetectable, see Fig. S7 and SI Appendix). Time-dependent sigmoidal kinetic profiles are associated with autocatalytic events (50) as in a dynamic homooligomerization constantly transferring folded monomers to a pool of unfolding-resistant or partially unfolding-resistant helical multimers. A detailed account of the development of the final form of the kinetic equations is provided in the SI Appendix; briefly, the unfolding kinetic constant was substituted by a sigmoidal function depending from the concentration of free folded monomers (as in autocatalysis) and thus from the dilution factor, the equations then solved numerically and plotted for comparison with the experimental data. It can be seen that, while lacking some quantitative insights (we cannot derive, for example, from the data in our hands, the average size of the multimers or the microscopic kinetic constants) the equations describe effectively the shape of the three curves and reproduce the trend observed with the change in the dilution factor.

We concluded that the intermittent contact between folded αSyn monomers, forming short-lived helical multimers, preserves the α-helicity of the unbound monomeric protein.

## Discussion

In this study we investigated the interplay of αSyn with 13:0 PC SUVs, binding partners characterized by the dependence of the protein’s membrane affinity on the structure of the lipid bilayers, akin of its interaction with lipid rafts (34, 37). By modulating αSyn’s binding, we were able to reconstitute a soluble helical species and investigate the nature of the folded conformer and its range of stability. αSyn retains most of its α-helical content after the affinity for lipid interfaces is “switched off” (Fig. 1B, Fig. S3B) and remains in a metastable folded conformation for an extended period of time, undergoing slow unfolding (Fig. 4D). The hysteresis in the helical fold is shown to be uncoupled from structural transitions of the lipid bilayer (Fig. 3) and several independent techniques prove that the folded conformers of αSyn are largely lipid-unbound (Fig. 4A-C, Fig. 5). Finally, the concentration-dependence of the unfolding rates and the kinetic analysis of the unfolding curves allow us to conclude that the helical species observed are stabilized by dynamic, cooperative multimerization (Fig. 5B-E). Lipid interfaces thus act as molecular chaperones for αSyn’s monomeric helical form and dynamic homo-oligomerization confers it its kinetical stability.

This phenomenon, previously unreported for IDPs, could open a new chapter in the already multifaceted behavior of this protein family. While folding of IDPs upon binding to ligands (either small molecules, nucleic acids or other proteins) has been long known and characterized in a wide number of cases (51–53), there is no previous report describing folding assisted by intermittent contact with a cofactor. In fact, IDPs for which binding and folding are coupled phenomena have binding enthalpies high enough to balance the entropically disadvantageous process of folding (52). The folding process described in this report seems to lie at the crossroads of the two mechanistic extremes of IDPs’ binding and folding models. αSyn’s folding during the downscan branch of the curve, could be described as “induced folding” (53) as it is the contact with the curved bilayers, starting with the insertion of the N-terminus in the lipid packing defects (54), that causes the protein to switch its conformation. On the other hand, the dynamic homo-oligomerization that takes place between the folded monomers is a “conformational selection” mechanism (53) as the helical amphipathic conformers engage in the intermittent contact that leads to their stabilization, as shown by the CD-followed unfolding curves and the related kinetic analysis (Fig. 5 and SI Appendix). The slow unfolding of helical αSyn (Fig. 4D) is thus the result of the competition between the irreversible unfolding of αSyn and the reversible oligomerization of folded monomers. A monomeric α-helical species of αSyn, given its extremely long half-life and its propensity for oligomerization, albeit metastable, is also likely to be the key intermediate in the folding mechanism of physiologic α-helical multimers (17, 23, 24, 27, 29, 55, 56). αSyn would then join several other IDPs that stably fold only upon homo-oligomerization (57–60), with a mechanism that could be described as conformational selection through a liposomal chaperone (Fig. 6). It is critical to stress how the absence, in our reconstituted αSyn sample, of stable homo-oligomers and the intermittent nature of the self-association are testaments to the reductionist nature of our experimental design. While lipid interfaces almost certainly play a central role in the refolding and assembly of physiological multimers, one or more key cofactors present in the eukaryotic cell cytosol (either molecular crowding, metal ions, other proteins or soluble lipids/fatty acids) appear to be necessary to stabilize these assemblies (30). αSyn multimers have been reported to localize at the synaptic bouton, around vesicles or vesicular clusters, but no consensus exists regarding their soluble or membrane-bound nature (17, 55). The apparent importance of transient membrane binding in αSyn’s conformational homeostasis might explain the difficulties encountered trying to answer this question, as these complex equilibria would not allow efficient trapping and localization of helical multimers without the employment of elaborate purification protocols (23) or techniques, such as NMR, that allow the detection of short-lived and kinetically-trapped species (24). Some groups have conducted structural studies of αSyn via NMR in a cytoplasmic environment (61, 62). Nevertheless, it remains difficult to obtain clear-cut answers given the close resemblance between the shifts of monomeric αSyn and its tetrameric form (61), the requirement that these in-cell experiments be performed in eukaryotic systems given the absence of potentially critical factors for folding and oligomerization in prokaryotic systems (30, 31), and the low sensitivity of NMR, especially for protein complexes and membrane-interacting proteins (63). The conclusions of the present work also concur with past reports arguing for the possibility of a more dynamic interaction of αSyn with curved membranes. While αSyn’s affinity for charged curved lipid interfaces has long been known (10) and its localization to the synapse has been established since its discovery (64), it remains consistently purified as a predominantly cytosolic protein (34, 65). Recently, a combination of solution and solid-state NMR measurements showed that, while its N-terminus is membrane-bound, the central (NAC) region of αSyn exists as an ensemble of transiently folded conformations closely associated with (but not bound to) membranes mimicking synaptic vesicles (66). Furthermore, fluorescence recovery after photobleaching (FRAP) experiments performed on primary neuronal cultures have found αSyn’s mobility to be lower than soluble proteins, such as GFP, but higher than synaptic vesicle proteins such as synapsin and SV2, suggesting a loose association with vesicles (35), closely resembling the transient interaction with vesicles on which our refolding paradigm relies. Similar results have also been obtained in murine models via multiphoton imaging (67). Together, these reports suggest a possible mechanism underlying aggregation and toxicity in synucleinopathies. αSyn’s homeostasis in the cell appears to regulate the protein’s distribution between two pools: one of them aggregation-prone, soluble, and unfolded; the other α-helical and aggregation-resistant (either membrane-bound, membrane-associated or soluble (23, 55, 68)). The relative amounts of these two populations of protein may correlate with synaptic activity, as suggested by the dispersion of αSyn in response to continuous stimulation (35) (Fig. 6). Imbalances in the relative amounts of protein in these two pools might then lead to nucleation events and the inception of aggregation (31). While some groups have advanced the hypothesis that membrane-binding actually favors the early events of aggregation (44), the transience of αSyn’s interaction reported here might prevent the increase in local concentration triggering nucleation events. The key to this modulation might reside, as the same authors recently suggested, in αSyn’s varying affinity for lipid interfaces and membrane microheterogeneities (69), or in cellular factors that regulate its membrane association (70, 71). This model could potentially reconcile disparate PD-associated clinical findings, from the disease risk conferred by altered brain lipid composition due to GBA mutations (22), to the abnormal lipid binding and tetramer destabilization of αSyn carrying fPD-causing missense mutations (48, 72–74).

**Fig. 6.**
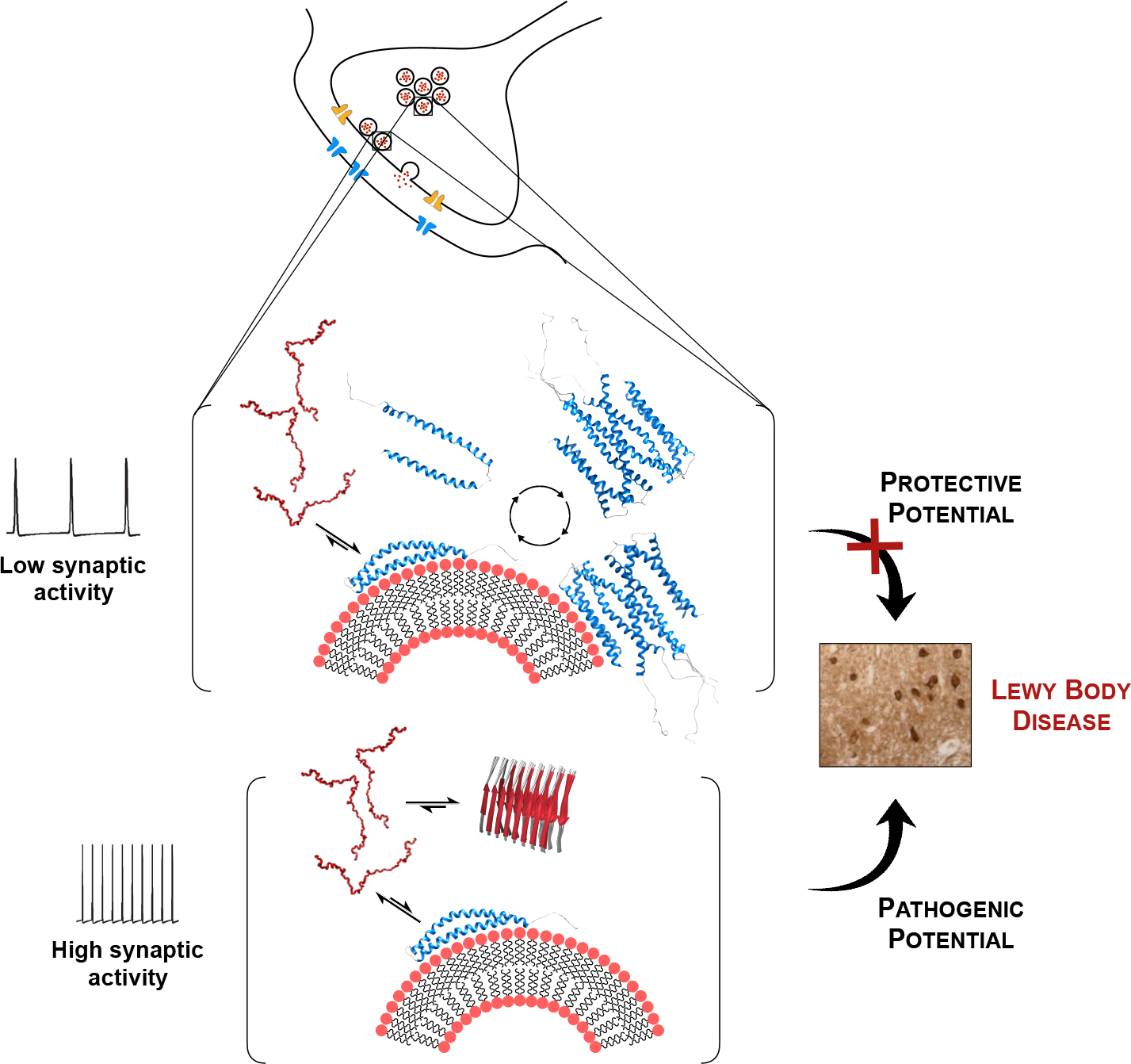
Model of helical αSyn monomers as critical intermediates in physiology and pathology. Normally, through its interaction with membranes, αSyn could retain its helical fold when unbound from the lipid interfaces and undergo homo-oligomerization into aggregation-resistant helical multimers (23) favored by its amphiphilic nature *(top)*. High synaptic activity has been shown to cause “dispersion” of αSyn, probably due to the constant membrane remodeling (35), and would thus drive a drastic decrease in the membrane-bound and membrane-associated protein pool, either monomeric or multimeric *(bottom)* The consequent dyshomeostasis would enlarge the pool of aggregation-prone unfolded monomeric αSyn. Similar effects could be caused by missense mutations, multiplications of the SNCA locus and lipid imbalance (31).

## Acknowledgments

We thank Kelly Arnett and the Center for Macromolecular Interactions at the Harvard Medical School Department of Biological Chemistry and Molecular Pharmacology for assistance with CD and ITC measurements and many helpful discussions; Simon Jenni, Yoana Dimitrova and the Harrison laboratory at the Harvard Medical School Department of Biological Chemistry and Molecular Pharmacology for assistance with DLS measurements; Maria Ericsson and the Harvard Medical School Electron Microscopy Facility for assistance with transmission electron microscopy; Alessandro Achille for his invaluable support in the development of the kinetic analysis. We also gratefully acknowledge Debby Pheasant and the MIT Biophysical Instrumentation Facility for the Study of Complex Macromolecular Systems (NSF-0070319) for the use of instruments and assistance with DSC measurements. Finally, we would like to thank Adam Cantlon, Erica Grignaschi, Ulf Dettmer, Silke Nuber, and all the other members of the Bartels, Dettmer and Selkoe laboratories at the Ann Romney Center for Neurologic Diseases for many helpful discussions. This work was supported by the Brigham and Women’s Parkinson’s Disease Research Fund, the American Parkinson’s Disease Association Research Grant and the MARDC Pilot Grant (to T.B.).

